# Visual perceptual learning of a primitive feature in human V1/V2 as a result of unconscious processing, revealed by Decoded fMRI Neurofeedback (DecNef)

**DOI:** 10.1101/2020.11.30.405209

**Authors:** Zhiyan Wang, Masako Tamaki, Kazuhisa Shibata, Michael S. Worden, Takashi Yamada, Mitsuo Kawato, Yuka Sasaki, Takeo Watanabe

## Abstract

While numerous studies have shown that visual perceptual learning (VPL) occurs as a result of exposure to a visual feature in a task-irrelevant manner, the underlying neural mechanism is poorly understood. In a previous psychophysical study, subjects were repeatedly exposed to a task-irrelevant global motion display that induced the perception of not only the local motions but also a global motion moving in the direction of the spatiotemporal average of the local motion vectors. As a result, subjects enhanced their sensitivity only to the local moving directions, suggesting that early visual areas (V1/V2) that process local motions are involved in task-irrelevant VPL. However, this hypothesis has never been examined by directly examining the involvement of early visual areas (V1/V2). Here, we employed a decoded neurofeedback technique (DecNef) using functional magnetic resonance imaging. During the DecNef training, subjects were trained to induce the activity patterns in V1/V2 that were similar to those evoked by the actual presentation of the global motion display. The DecNef training was conducted with neither the actual presentation of the display nor the subjects’ awareness of the purpose of the experiment. As a result, subjects increased the sensitivity to the local motion directions but not specifically to the global motion direction. The training effect was strictly confined to V1/V2. Moreover, subjects reported that they neither perceived nor imagined any motion during the DecNef training. These results together suggest that that V1/V2 are sufficient for exposure-based task-irrelevant VPL to occur unconsciously.

**Significance Statement:** While numerous studies have shown that visual perceptual learning (VPL) occurs as a result of exposure to a visual feature in a task-irrelevant manner, the underlying neural mechanism is poorly understood. Previous psychophysical experiments suggest that early visual areas (V1/V2) are involved in task-irrelevant VPL. However, this hypothesis has never been examined by directly examining the involvement of early visual areas (V1/V2). Here, using decoded fMRI neurofeedback, the activity patterns similar to those evoked by the presentation of a complex motion display were repeatedly induced only in early visual areas. The training sensitized only the local motion directions and not the global motion direction, suggesting that V1/V2 are involved in task-irrelevant VPL.

## Introduction

Visual perceptual learning (VPL) refers to the long-term performance enhancement resulting from visual experiences (Sagi, 2011; Watanabe & Sasaki, 2015). VPL can occur from mere exposure to a visual feature (Arsenault & Vanduffel, 2019; Galliussi, Grzeczkowski, Gerbino, Herzog, & Bernardis, 2018; Gutnisky, Hansen, Iliescu, & Dragoi, 2009; Lorenzino & Caudek, 2015; Pascucci, Mastropasqua, & Turatto, 2015; Protopapas et al., 2017; Rosenthal & Humphreys, 2010; Seitz & Watanabe, 2003; Watanabe et al., 2002; Watanabe, Nanez, & Sasaki, 2001). Despite the multitude of studies on this phenomenon, the underlying mechanism of exposure-based task-irrelevant VPL remains unclear.

Psychophysical experiments have been conducted to infer the mechanism of exposure-based VPL. In Watanabe et al. (2002), the researchers peripherally presented a motion display, termed the “Sekuler display”, which consisted of dots moving randomly within a certain range of directions as a task-irrelevant stimulus while subjects were asked to conduct another task presented at the center of the display. This display induced the perception of not only the local dots’ motions but also the global motion corresponding to the spatiotemporal average of the local motion vectors (Williams & Sekuler, 1984). After repetitive exposures to the Sekuler display as task-irrelevant stimuli, the sensitivity within the local dots’ direction range enhanced. In another condition where a global motion discrimination task was trained in the Sekuler display, in the earlier phases of training, the sensitivity within the local dots’ direction ranges enhanced. However, in later phases of training, sensitivity enhancement was observed in the global motion direction only if a task was conducted on the global motion. Given that V1 is the first cortical area to respond to local motion directions and V3A is the first area to respond to global motion (Koyama et al., 2005), these findings suggest that in early phases of training, VPL occurs on local features by passive exposure, and in later phases, VPL develops based on a task-relevant global feature that occurs in higher-level stages of visual processing.

Harris, Gliksberg, and Sagi (2012) conducted training on orientation discrimination of popped-out texture elements from a background texture. As in previous studies (Karni & Sagi, 1991, 1993), VPL of this task was found to be specific for the location where the target was presented. However, training of another condition in which target presentation was interleaved by task-irrelevant counterorientation trials led to transfer of the VPL of the task to untrained locations. Given that such trials in a similar setting were found to abolish an aftereffect that is involved in a lower-level, location-specific stage (Greenlee & Magnussen, 1988), the location nonspecificity in this condition may be due to the elimination of a lower-level location-specific plasticity in VPL. The results of these studies lead to the hypothesis that VPL includes changes in the processing of local feature in early visual cortical areas in a passive manner, irrespective of whether the feature is task-relevant.

To test this hypothesis, we conducted experiments with decoded fMRI neurofeedback (DecNef) training. A number of studies have found that DecNef training repetitively induces patterns similar to those evoked by a real stimulus without presentation of the stimulus, enhancing performance on the stimulus (Amano, Shibata, Kawato, Sasaki, & Watanabe, 2016; Shibata, Watanabe, Sasaki, & Kawato, 2011; Watanabe, Sasaki, Shibata, & Kawato, 2017).

We found that repetitive inductions of activity patterns in V1/V2 by DecNef similar to those evoked by the Sekuler display resulted in performance enhancement of the local motion directions and not specifically of the global motion direction to which V3A is the earliest area to respond (Koyama et al., 2005). A leak analysis of BOLD activity showed that only the V1/V2 activity patterns were correlated with the neurofeedback training. The subjects’ reports showed no evidence that they had perceived or imagined any motion during the DecNef training. These results are in accordance with the hypotheses that VPL includes changes in the processing of local features in V1/V2 in a passive manner.

## Materials and Methods

### Subjects

Fourteen subjects (aged 18-30) with normal or corrected-to-normal vision participated in the study. Subjects had no prior experience participating in visual training experiments and were naïve or had little knowledge about brain anatomy. Eight subjects participated in the neurofeedback experiment and six subjects participated in the control experiment. All subjects gave written consent to the study, which was approved by the Institutional Review Board of Brown University. The number of subjects was comparable to that of previous neurofeedback studies examining visual perception (Amano et al., 2016; Scharnowski, Hutton, Josephs, Weiskopf, & Rees, 2012; Shibata et al., 2011).

### Neurofeedback Experiment

#### Experiment Timeline

The neurofeedback experiment consisted of 4 stages as shown in Figure 1A: the pretest stage (1 session), fMRI motion decoder construction stage (1 session), fMRI neurofeedback training stage (3 sessions), and posttest stage (1 session). Each session was conducted at least 24 hours apart. See below for each stage.

**Figure 1.**
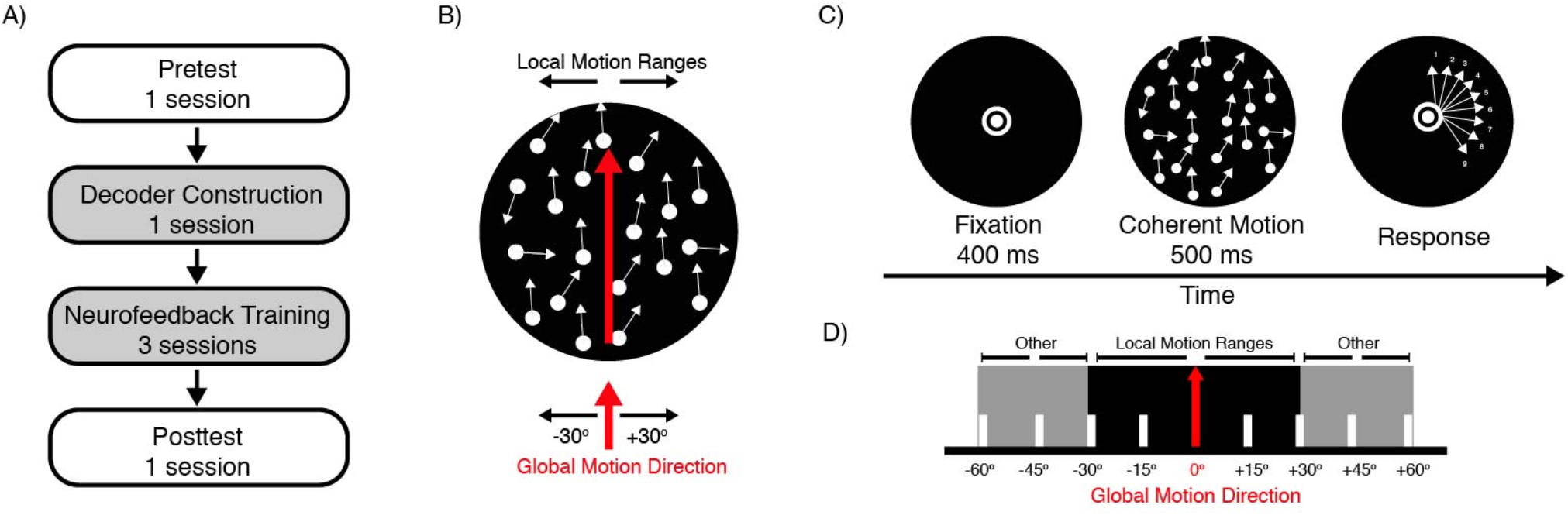
Experiment design and stimuli. A) Experiment timeline. Decoder construction and neurofeedback training stage were performed inside MRI scanner (gray background). Pre- and posttest stages were performed psychophysically. B) Global motion display. Local motions were within +/- 30° range of the global motion direction. C) Paradigm for pre- and posttests. D) Illustration of the 9 motion directions during pre- and posttests. The motion directions covered the global motion direction, the local motion ranges (from −30° to +30° relative to the global motion direction) and other motion directions outside the range of the global motion display (from −60° to −30° and from +30° to +60° relative to the global motion direction).

#### Statistical Analysis

The normality of the data was checked with the Shapiro-Wilk test. All datasets were normally distributed. Therefore, t-tests were applied to determine statistical significance in the behavior measures. Effect sizes were also calculated accompanying the statistical significance. When necessary, Bonferroni correction was applied to correct for multiple comparisons. For leak analysis, permutation tests were performed to acquire significance (see *Permutation test* for details). All of the statistical analyses were performed using MATLAB (MathWorks, Natick, MA). Shapiro-Wilk tests were performed with MATLAB functions developed by Öner and Deveci Kocakoç (2017).

#### Motion Stimuli

There were two types of motion used. One was a coherent motion (Braddick et al., 2001; Salzman, Murasugi, Britten, & Newsome, 1992). The other was a “Sekuler display” with additional noise overlaid (Williams & Sekuler, 1984).

During the pre- and posttest stages, a 10% coherent motion display was presented. The display consisted of 70 dots moving within an aperture with a radius of 4.5°. For each frame, 10% of the dots (coherent dots) moved coherently in a predetermined direction. The coherent dots moved at a speed of 7.1°/sec. The other dots were randomly placed at different locations from frame to frame. Each frame was presented for ~16.7 ms. See Pre- and Posttest stage below for more details.

During the fMRI motion decoder construction stage, a Sekular display was presented as shown in Figure 1B. The motion display consists of local motions moving spatiotemporally and randomly within a certain range (−30° – 30° relative to the global motion direction). The display induces a global motion perception in the direction of the spatiotemporal average of the local motion signals (Koyama et al., 2005; Williams & Sekuler, 1984). For the global motion stimuli, 35 signal dots and 65 noise dots moved within an aperture (4.5° radius) presented at the center of the display. For each frame, each signal dot moved in a random direction uniformly distributed within the range from a predetermined global motion direction, defined as the mean of the distribution of the local motion signals. Each noise dot moved randomly without any direction restriction. The subjects perceived both the local motion of each moving dot as well as the global motion. Each frame was presented for 50 ms. Two ranges of global motion displays were used. The global motion direction in one range was rotated 120° from that in the other range so that there was no overlap between the two ranges of global motion display. One range of global motion direction corresponded to the trained motion range for neurofeedback. The other range was used for control (untrained). See *fMRI Motion Decoder Construction* for more details.

The background was black except for a white bull’s eye on a gray disc with a radius of 0.75° presented at the center of the display in both the fMRI motion decoder construction stage and the testing stages.

#### Apparatus

The visual stimuli used during the behavioral testing sessions were presented on an LCD monitor (1024 × 768 resolution, 60 Hz refresh rate), whereas those used during the MRI sessions were presented on an LCD display (BOLDscreen 32 LCD for fMRI, Cambridge Research Systems, 1920 × 1080 resolution, 120 Hz refresh rate). The visual stimuli used during the behavior testing sessions and decoder construction sessions were controlled via a Mac OS computer and Psychtoolbox (Brainard, 1997). The visual stimuli used during the neurofeedback training sessions were controlled via a Windows 7 computer and Psychtoolbox (Brainard, 1997).

#### Pre- and Posttests

The purpose of the pre- and posttest stages was to measure subjects’ sensitivity to each motion direction within or beyond the local motion direction range in the Sekuler display. The pre- and posttest stages were conducted in a dimly lighted room. A total of 18 motion directions was tested. The sensitivity of each motion direction was measured by performing 20 trials of motion discrimination tasks in each of the pretest and posttest stages. In each test stage, there were 2 blocks; each block presented either the trained or the untrained global motion direction as well as other 8 motion directions (a total of 9 motion directions each block). Within each block, the order of the presentation of the 9 motion directions was pseudo-randomized. In Figure 1D, we illustrated the 9 directions that included one of the global motion displays. The range of the 9 motion directions covered the global motion direction (shown as 0°), the local motion directions (from −30° to +30° relative to the global motion direction) as well as other motion directions (from −60° to −30°, and from +30° to +60° relative to the global motion direction) outside the local motion directions. Two neighboring directions were 15° apart from each other.

In one trial, the subjects performed a motion discrimination task with 10% coherent motion stimuli (shown in Figure 1C). At the beginning of each trial, a fixation point was presented for 400 ms. A coherent motion display with 10% coherence was presented for 500 ms. The stimulus was followed by a response screen with 9 arrows and corresponding keyboard buttons indicating the possible motion directions. In the response screen, the range of 9 arrows were fixed for each block. The subjects were asked to indicate which motion direction was presented by pressing the corresponding key on the keyboard.

For each direction the behavioral performance was characterized by a change between pre- and posttest in d-prime, which was calculated as *z* (Hit rate) – *z* (False alarm). The tuning function of the performance changes as a function of the motion direction was acquired by fitting a smooth spline with piecewise polynomials to the mean d-prime across subjects at each motion direction. The smoothing parameter was 0.95. The mean improvement in the each of trained and untrained local motion ranges was calculated by summing the d-prime for each direction across the trained / untrained local motion range per subject (including the global motion direction) and averaged across subjects.

#### fMRI Motion Decoder Construction

The fMRI motion decoder construction stage was conducted while the subjects stayed inside the scanner to obtain the blood-oxygen-level dependent (BOLD) signal patterns in V1/V2 (see below) produced in response to the two global motion directions.

Initially, the subjects’ structural anatomical brain images were acquired with Siemens’ AutoAlign function (see MRI parameters). AutoAlign ensured that the subsequent functional scans were automatically positioned on the same slice of the brain, which was aligned to the plane equivalent to the anterior commissure to posterior commissure (AC-PC) plane.

Then, the subjects’ BOLD signals were collected with decoder construction functional runs while they were passively exposed to the two global motion stimuli. The subjects were instructed to maintain fixation on a white bull’s eye on a gray disc while exposed to 240 trials of the global motion stimulus for a total of 10 runs. Each run lasted 300 sec in total and consisted of 24 trials of global motion stimuli (12 trials for each global motion direction). A 10-sec fixation period and a 2-sec fixation period were inserted before and after the trials, respectively. The initial fixation period was added to allow the magnetic field to reach a steady state. Each trial started with a 6-sec stimulus presentation period followed by a 6-sec fixation period. During the stimulus presentation period, one of the two global motion directions was presented for 6 sec, during which the color of the fixation point could change from white to green for 500 ms.

The fixation point changed color in 50% of the trials (12 trials in one run). In the remaining 50% of the trials, the color of the fixation point stayed white. The stimulus period was followed by the 6-sec fixation period, during which the fixation color stayed white. The fixation task was performed to ensure that subjects maintain their fixation during the decoder construction stage. The order of trials with and without color change were randomly determined for each subject. The subjects were asked to press a button with their index finger if they noticed a color change during the 6-sec fixation period.

Decoder construction runs were preprocessed with FreeSurfer (Fischl, 2012) software. Each run underwent 3-D motion correction to align all of the functional runs to the first of the decoder construction runs. The average motion for each subject across different runs was less than one voxel. Rigid body transformation was then performed individually to align the functional runs to the structural image. A gray matter mask was acquired for further analysis.

In the same session following the decoder construction, retinotopy mapping runs were performed with a standard retinotopic procedure to delineate the visual areas of each subject (Engel et al., 1994; Fize et al., 2003; Yotsumoto, Watanabe, & Sasaki, 2008). The retinotopic stimulus consisted of a colorful checkerboard that extended from 1° to 4.35°. The retinotopic stimuli occupied a smaller region than the global motion stimuli to avoid selecting voxels that corresponded to the edges of the stimuli. The checkerboard stimuli alternated between horizontal and vertical meridians as well as upper and lower visual fields. The subjects fixated on the fixation point and pressed a button when they noticed that it changed color. For each frame of stimulus presentation, there is a 0.01 probability that the color of the fixation point changed to red. Retinotopy mapping runs were preprocessed with FreeSurfer software. 3-D motion correction was performed with the first decoder construction run as the template. Brain mask and intensity normalization were performed. Rigid body transformation was performed to align the retinotopy runs with the structural image. V1/V2 were delineated by computing the contrast between the horizontal and vertical meridians and projected onto the inflated brain surface.

Next, the time-course of BOLD activations from the decoder construction runs were extracted from the voxels corresponding to the V1/V2 region defined in the retinotopy mapping with MATLAB. The functional data were then shifted by 6 secs to account for the hemodynamic delay. Detrending analysis was performed to remove linear trends from the functional runs, followed by z-score normalization. The data samples used in decoding for each trial were the averaged intensities of 3 functional volumes that corresponded to each 6-sec stimulus period. Thus, we acquired 240 data samples that corresponded to the 240 trials.

Finally, we obtained the decoder for the two global motions with sparse logistic regression (Miyawaki et al., 2008; Yamashita, Sato, Yoshioka, Tong, & Kamitani, 2008). The input to the algorithm was the 240 data samples in V1/V2. Sparse logistic regression automatically selected voxels relevant for the separation of the representation of the two global motion directions. The output from the calculation was a decoder that consisted of estimated weights for each selected voxel in V1/V2. Leave-one-out cross-validation was conducted separately. For each round of the cross-validation, 216 data samples from 9 runs were used to train the decoder with sparse logistic regression. The accuracy of the decoder was tested with the 24 samples from the remaining run. The decoder was used to predict the likelihood of each global motion direction and assign each direction to each test sample. The accuracy of the decoder’s prediction was then calculated by comparing the predicted direction with the true global motion direction that was presented. Thus, we acquired the test accuracy for each run in each cross-validation round, and the accuracy of the cross-validation was averaged across 10 cross-validation rounds.

#### fMRI Neurofeedback Training

The decoder constructed during the decoder construction session was applied to the neurofeedback training sessions. The 8^th^ volume image from the first decoder construction run was also obtained as a template image for real-time evaluation of the quality (e.g. motion artifact) of the functional runs (see below). One of the two global motion displays was pseudorandomly selected as the trained global motion display for each subject. Initially, a structural scan with AutoAlign was performed to automatically place the slice on the same plane as during the decoder construction sessions.

Then, we performed a number of induction runs in which subjects attempted to induce the trained global motion pattern in V1/V2 with neurofeedback without realizing they were being trained (see below). Each induction run lasted for 330 sec, starting with a 30-sec fixation period followed by fifteen 20-sec trials, as shown in Figure 2. The subjects were instructed to fixate on the fixation point throughout the runs. Each trial started with a 6-sec induction period. During the induction period, a ‘+’ sign was presented at the center of the screen while the subjects were instructed to try to use the posterior part of their brain to make a later-presented green disc as large as possible. The induction period was followed by a 6-sec fixation period, during which a ‘-’ sign was presented at the center of the screen while the subjects were instructed to fixate only. The fixation period was added to accommodate the 6-sec hemodynamic response delay. The feedback period was presented afterwards for 2 sec. A green disc was presented at the screen center, with the size of the disc being proportional to the likelihood of the trained global motion direction based on the current BOLD activation in V1/V2. There was a 5-deg gray boundary for the disc. Each trial ended with a 6-sec intertrial interval, while a ‘=’ sign was shown. Subjects were instructed to relax but maintain their fixation during the “=” period.

**Figure 2.**
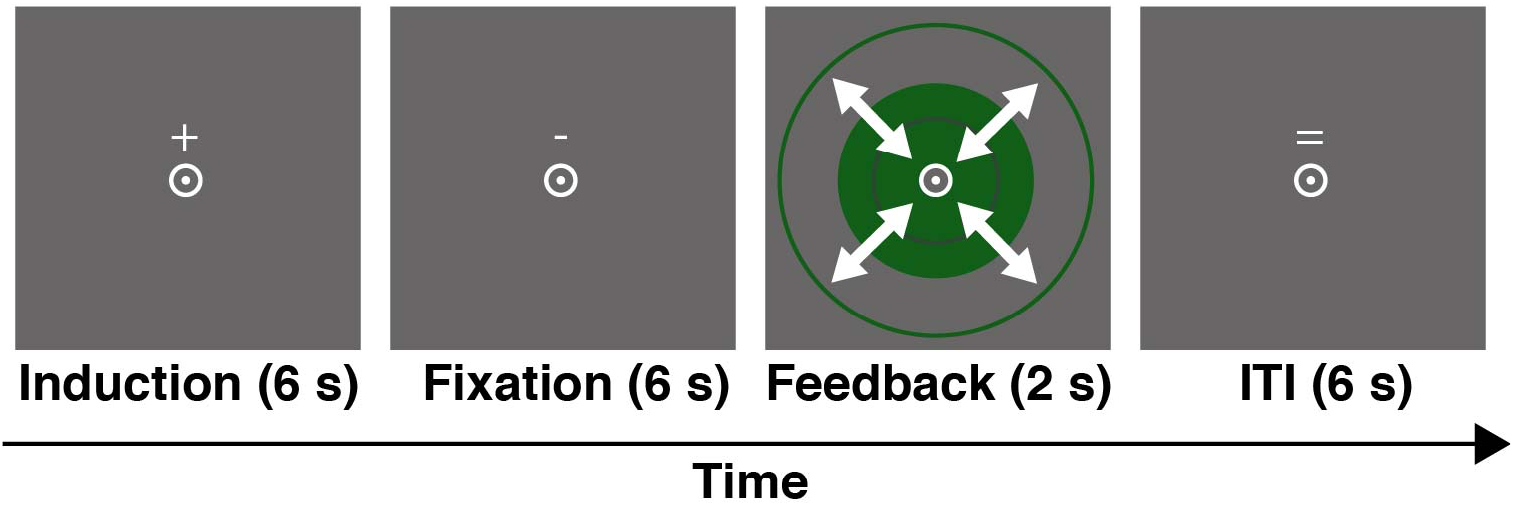
One trial in an induction run. A ‘+’ indicated the induction period, during which subjects tried to engage their brain activities to make a subsequently presented green disc larger. ‘-’ and ‘=’ indicated fixation and ITI periods, respectively. Feedback was given to the subjects for 2 seconds by the disk size. Subjects were instructed to maintain fixation throughout the run.

Each volume was transferred in real time from the scanner to a remote computer in which the real-time BOLD activations were calculated with MATLAB and SPM. The computation was performed as follows. The functional volumes that corresponded to the induction period were 3D motion corrected in real time with SPM. The spatial correlation was calculated between the current volume and the template acquired from the decoder construction session. The correlation score was only shown to the experimenter, who was outside the scanner room. A correlation less than 0.85 indicated a significant movement of the subjects in the scanner, which could lead to an inaccurate selection of voxels. Thus, the run was terminated if such a correlation was observed, and a structural scan with AutoAlign was performed for accurate functional slice placement. Later, the BOLD activation time course was selected from the voxels that corresponded to the selected voxels for decoding in V1/V2. Then, detrend analysis was performed to remove the linear trend from the time course of the BOLD activation. Following linear trend removal, z-score normalization was performed for each of the voxels to the 10 sec - 30 sec functional volumes that were acquired from the onset of the run. Next, the intensities of the functional volumes that corresponded to the 6-sec induction period were averaged to create the input data sample to sparse logistic regression. Finally, the likelihood of the current induction period was calculated with the data sample and the precalculated weights during the decoder construction stage. The likelihood ranged from 0% to 100% and was reflected proportionally by the size of the green disc.

Subjects were allowed to take a break upon request. The mean (+/- SD) number of induction runs for each neurofeedback training session was 10.63 +/- 1.5 runs. After completion of each of the neurofeedback training session, the subjects were asked to freely report what strategies they took and/or what they had in mind during each of the three sessions. The results of their strategies are shown in Table 1. No reported strategy from any subject was related to motion.

**Table 1.** Reported strategies during the neurofeedback training stage

|  | Session 1 | Session 2 | Session 3 |
| --- | --- | --- | --- |
| Subject 1 | Visualizing green<br>disk | Visualizing green<br>disk | Visualizing green<br>disk |
| Subject 2 | Visualizing huge<br>green disk | Visualizing disk<br>getting larger | Visualizing disk<br>getting larger |
| Subject 3 | Thinking of green | Visualizing large<br>green circle | Thinking of green |
| Subject 4 | Emotional argument,<br>random feelings of<br>life, contemplations | Emotional argument,<br>sad stories | Contemplations,<br>imagining some<br>details of landscape<br>pictures |
| Subject 5 | Happy memories and events | Job and family members | Music and dancing |
| Subject 6 | Visualizing a circle getting larger | Visual experiences and imagining the circle getting larger | Visualizing objects, writing letters, etc. |
| Subject 7 | Peaceful images, landscapes involving green such as ponds, grass. | Movie scene from ‘Shining’, underwater scenes with algae | Maintaining fixation and imaging scenes with green elements. |
| Subject 8 | Musical notes and philosophical questions | Musical notes and philosophical questions | Musical notes and philosophical questions |

#### MRI Parameters

All of the subjects were scanned in a 3 Tesla Siemens PRISMA scanner with a 64-channel head coil. Each fMRI session, a T1-weighted MPRAGE sequence (256 slices, voxel = 1 mm * 1 mm * 1 mm, TR = 1980 ms, TE = 3 ms, flip angle = 9°), was acquired with AAScout AutoAlign (160 slices, voxel = 1.625 mm * 1.625 mm * 1.6 mm, 0 mm slice gap, TR = 3 ms, TE = 1.37 ms, flip angle = 8°) to ensure accurate placement of functional slices on the same location across sessions. For the retinotopy, decoder construction and induction runs, a gradient EPI sequence was acquired with 33 continuous slices (TR = 2 s, TE = 30 ms, flip angle = 90°, voxel = 3 mm * 3 mm * 3 mm) oriented parallel to the AC-PC plane placed by AutoAlign to cover the whole brain.

### Off-line Leak Analyses

We performed off-line leak analyses for the following reasons. Even if our fMRI neurofeedback training successfully induced the target motion-related activity pattern in V1/V2 and this resulted in VPL performance changes, it would not unequivocally indicate that the behavioral changes are explained merely by the induced activation pattern in V1/V2. It is possible that, in parallel with the specific activation pattern in V1/V2, similar activation patterns occurred in some other brain areas during the induction period of the neurofeedback training and contributed to the behavioral changes. Therefore, we tested whether the target local motion-related activity pattern in V1/V2 leaked out of the V1/V2 to other brain regions that are relevant for motion processing. To this end, we used the leak analysis, termed a region of interest (ROI)-based method as used in the previous studies (Shibata, Watanabe, Kawato, & Sasaki, 2016; Shibata et al., 2011). In the ROI-based method, we used anatomically delineated three motion-related ROIs in addition to V1/V2 and compared the amounts of leakage in these four regions (the V1/V2 and the other three regions). The details of the leak analysis are described in the following sections.

We used the ROIs of V1/V2 (the target area during neurofeedback training stage), V3A, the middle temporal area (MT), and the medial superior temporal area (MST) for this analysis because these regions are largely motion-related. While V1/V2 was delineated by the retinotopy mapping, V3A, MT, and MST were defined by using the parcellation of the human cerebral cortex in the FreeSurfer template brain (Glasser et al., 2016). After defining ROIs, we reconstructed the likelihood of the target motion-related activity in V1/V2 that was obtained online (in real time) using a sparse linear regression algorithm (Toda, Imamizu, Kawato, & Sato, 2011) based on activation patterns measured in each of the aforementioned four ROIs (i.e. V1/V2, V3A, MT, and MST) during the induction period of the neurofeedback training. Note that the reconstructed values from V1/V2 was not necessarily the same as the likelihood of the target motion-related activity pattern in V1/V2 obtained on-line because the preprocessing pipelines of fMRI signals were different between on-line and off-line procedures (see *Preprocessing for Leak Analysis*). Moreover, the likelihood we obtained on-line was the result of a non-linear logistic function (0% to 100%). Before we conducted the sparse linear regression, we transformed the likelihood of the target motion-related activity with a hyperbolic tangent function. We trained the sparse linear regression (Shibata et al., 2016; Shibata et al., 2011) using data from two of the neurofeedback training sessions and then calculated the reconstructed value of the rest of the neurofeedback training session in each ROI. Then we estimated the reconstruction performance of each ROI as a Fisher-transformed correlation coefficient between the reconstructed value and the likelihood of the target motion-related activity pattern in V1/V2 we obtained on-line. After the estimation of the performance, we conducted permutation tests (see *Permutation Test*) to evaluate the statistical significance of the estimated Fisher-transformed correlation coefficient for each ROI.

#### Preprocessing for Leak Analysis

The functional runs from the neurofeedback stage were preprocessed with FreeSurfer and MATLAB. All of the volumes underwent 3D motion correction with the first run of the decoder construction stage as the template volume. A gray matter mask of the whole brain was made. The BOLD activation time course was extracted from the voxels that corresponded to the whole-brain gray matter mask. The time course was shifted by 6-sec to account for the hemodynamic delay. Linear trends were removed, and z-score normalization was applied to the time course using the initial 10-sec – 30-sec of the time course. For the neurofeedback induction runs, data samples for the leak analysis were acquired by averaging the 6 sec induction period for each trial.

#### Permutation Test

For the correlation coefficients that were calculated with the leak analysis, we conducted a permutation test to acquire proper statistics. First, we permutated the order of the predicted probabilities from each of the 4 control ROIs 1000 times and obtained 1000 correlation coefficients between the reconstructed performance in the 4 ROIs and the likelihood of the target motion in V1/V2 which we acquired on-line. Second, we obtained the distribution of the correlation coefficients and tested whether the reconstructed performance acquired with leak analysis was among the top 5% of the permutation distribution. A correlation coefficient was regarded as significant if it ranked among the top 5% of the distribution. Third, we computed the z-score for each control ROI by comparing the acquired correlation coefficient with the permutation distribution for between-subject comparisons.

### Control Experiment

A behavioral control experiment was conducted outside of the scanner in a similar manner as in the neurofeedback training experiment but without induction training. The control experiment consisted of pre- and posttest sessions and a decoder exposure session. The procedures of the pre- and posttest sessions were the same as those in the neurofeedback experiment. Subjects were tested on 18 motion directions with 10% coherence. During the decoder exposure session, the subjects were presented with the same global motion stimuli and fixation task as in the decoder construction session except that the subjects were asked to respond to the fixation task by pressing a keyboard key. The posttest session was conducted at least 3 days apart from the decoder exposure session to ensure that the time courses of testing and training were as similar to those of the neurofeedback training sessions as possible.

### Code and Data availability

All data and customized code for analyzing behavior and fMRI results are available upon request.

## Results

### Neurofeedback Experiment

The neurofeedback experiment consisted of four stages: the pretest stage, fMRI motion decoder construction stage, fMRI neurofeedback training stage and posttest stage.

During fMRI decoder construction session, a decoder was constructed for individually defined V1/V2 to distinguish the BOLD activation patterns that corresponded to the two global motion stimuli with different global motion directions. The mean accuracy of the decoder across subjects was found to be significantly above the chance level [t(7) = 2.38, p = 0.04, Cohen’s d = 0.842] using cross-validation. One of the two global motion stimuli was selected as the trained pattern. During the fMRI neurofeedback training sessions, the subjects were instructed to induce the trained global motion pattern with an implicit feedback method (see **Materials and Methods** for details). The scores of neurofeedback training indicated the similarity between the subjects’ brain activation pattern and the trained global motion pattern. We measured the improvement in neurofeedback score as the change between the third and the first neurofeedback training session. The scores improved across the 3 neurofeedback training sessions. The mean improvement of neurofeedback training scores across subjects was 5.41 +/- 2.11 (SEM) and was significantly above zero [t(7) = 2.56, p=0.038, Cohen’s d = 0.905]. During the pretest and posttest stages, the subjects performed motion discrimination tasks for 9 motion directions surrounding each of the 2 global motion directions covering the local motion ranges (as in Figure 3). Each subject’s performance improvement in the motion discrimination tasks was characterized as an improvement in d-prime (Posttest d-prime – Pretest d-prime). A smooth spline curve was fitted for a visualization purpose to the mean d-prime improvement across the trained local motion directions and the untrained local motion directions, separately, as shown in Figure 3 and Figure 5.

**Figure 3.**
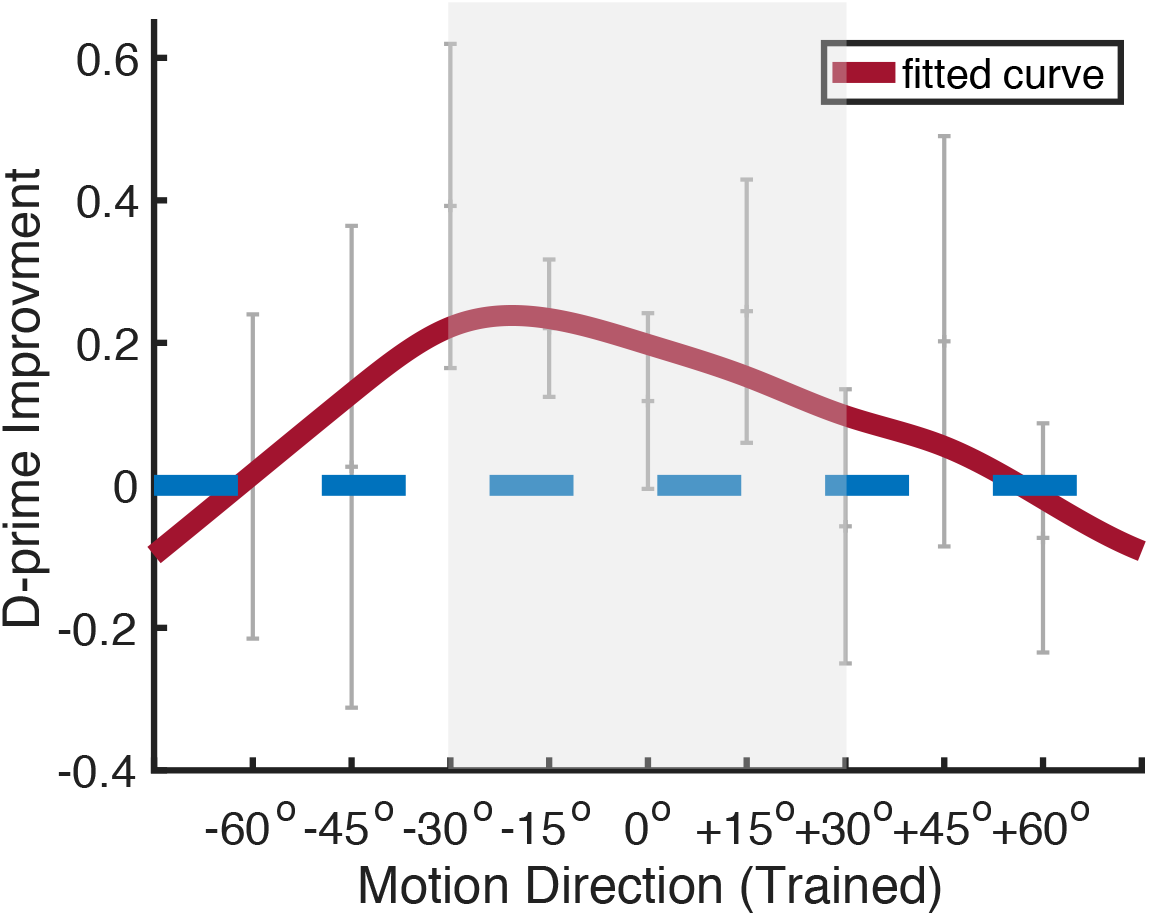
Mean (±SEM) d-prime improvement in the trained motion range. 0 deg represents the global motion direction. The red line represents the spline fitting of the data points. The shaded area represents the local motion direction range.

As shown in Figure 4, the d-prime improvement, summed across all local motion direction for each subject (see **Material and Methods** for detail), was significantly greater than zero [t(7) = 2.57, p = 0.037, Cohen’s d=0.91]. In contrast, no significant improvement was obtained in the untrained local motion range [t(7) = −0.866, p = 0.415, Cohen’s d = −0.3062]. The improvement for both the trained global motion direction [t(7) = 0.9601, p = 0.369, Cohen’s d = 0.339] and the untrained global motion direction [t(7) = 0.046, p = 0.965, Cohen’s d = −0.016] was not significantly greater than zero (0° motion direction in Figure 3 and Figure 5).

**Figure 4.**
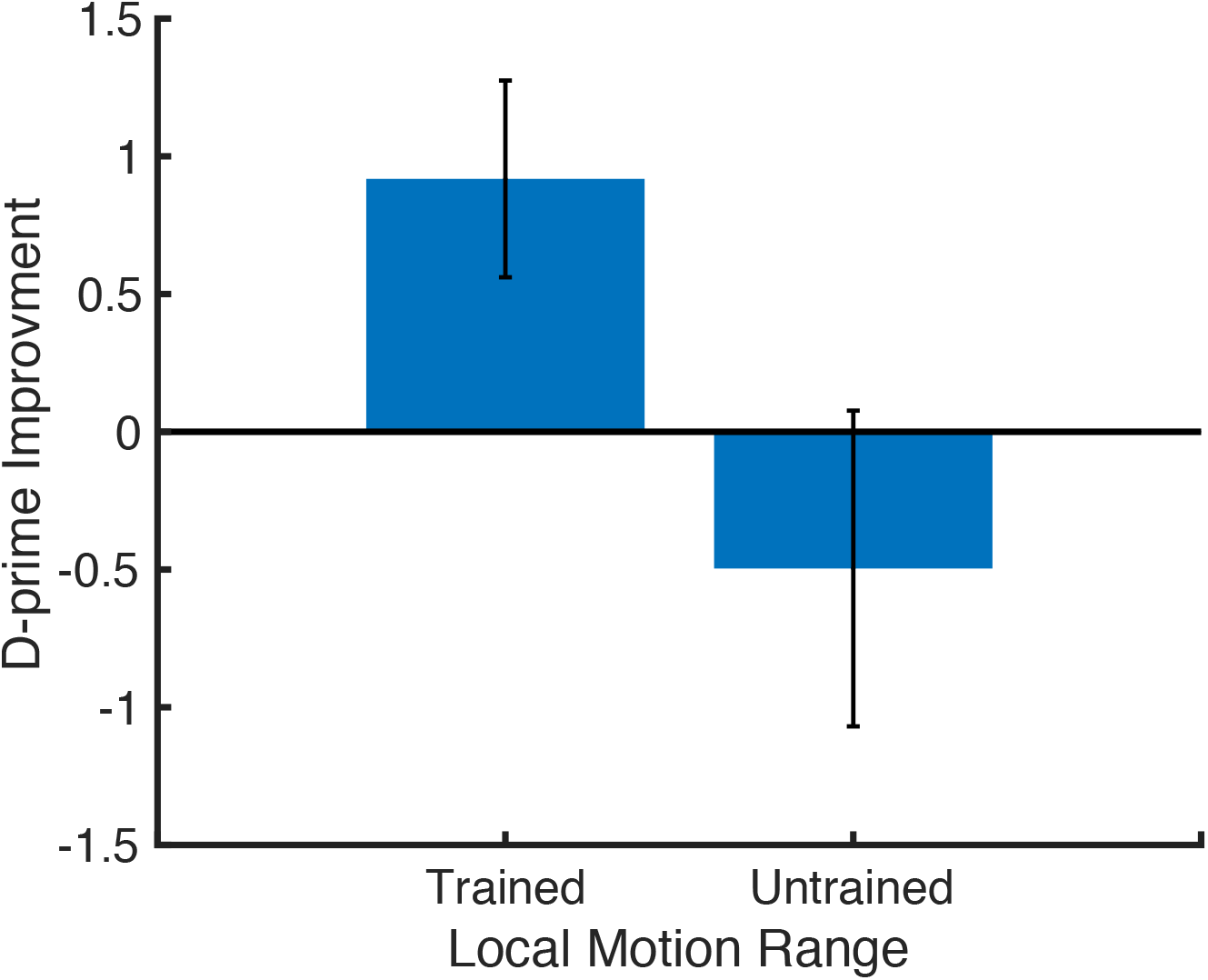
Mean (±SEM) improvements for trained vs untrained local motion ranges. There was significant improvement in trained local motion range. No significant improvement in the untrained local motion range was observed.

**Figure 5.**
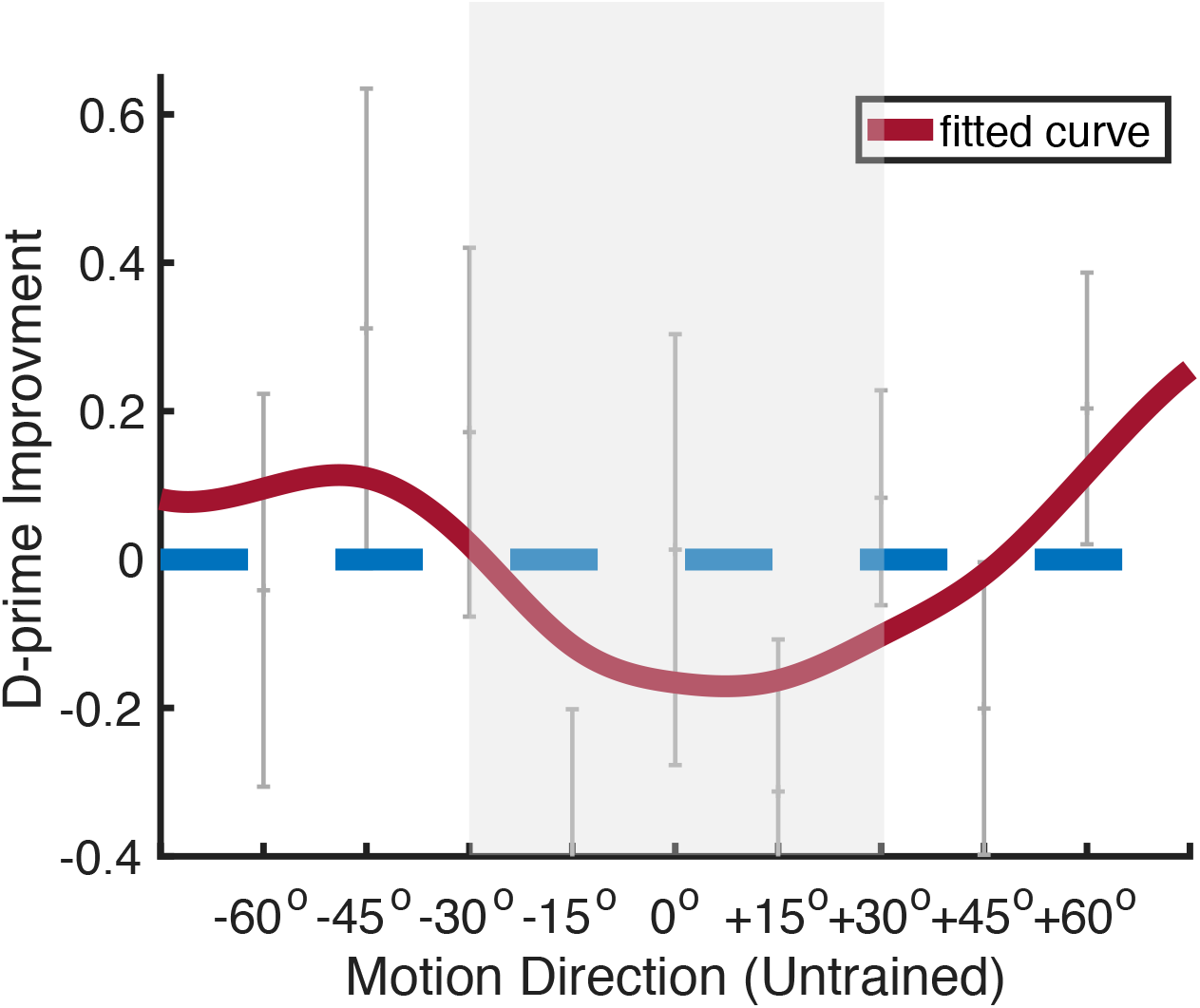
Mean (±SEM) d-prime improvement in the untrained motion range. 0 deg represents the global motion direction. The shaded area represents the local motion.

### ROI-based Leak Analysis

To ensure that the behavioral performance changes were driven by activity in V1/V2 alone and not in other visual areas or brain regions, we conducted an ROI-based leak analysis. We defined the motion-related brain regions (V1/V2, V3A, MT and MST) and measured the reconstructed probability of the trained motion pattern with sparse linear regression in each of the individual motion-related brain regions during the neurofeedback training stage. If V1/V2 alone induced the behavior performance during neurofeedback training, we would expect only V1/V2 to be capable of reconstructing the neurofeedback training scores, which should reflect the likelihood of the on-line brain activation pattern in V1/V2. If, on the other hand, other motion-related regions were also involved in the process, we would expect regions other than V1/V2 to have the capability of reconstructing the neurofeedback training scores. The reconstructed performance was defined as the Fisher-transformed correlation coefficient between the reconstructed probability and the estimated probability in V1/V2 from real-time experiments (see **Materials and Methods** for details). The reconstructed performance was significantly greater than zero for V1/V2 [z = 8.171, p=0.000]. The results also indicate that the correlation coefficient for any area other than V1/V2 was not significantly higher than zero [MT: z = 0.498, p = 0.31; MST: z = 0.569, p = 0.28; V3A: z = −0.421, p = 0.66] (Figure 6).

**Figure 6.**
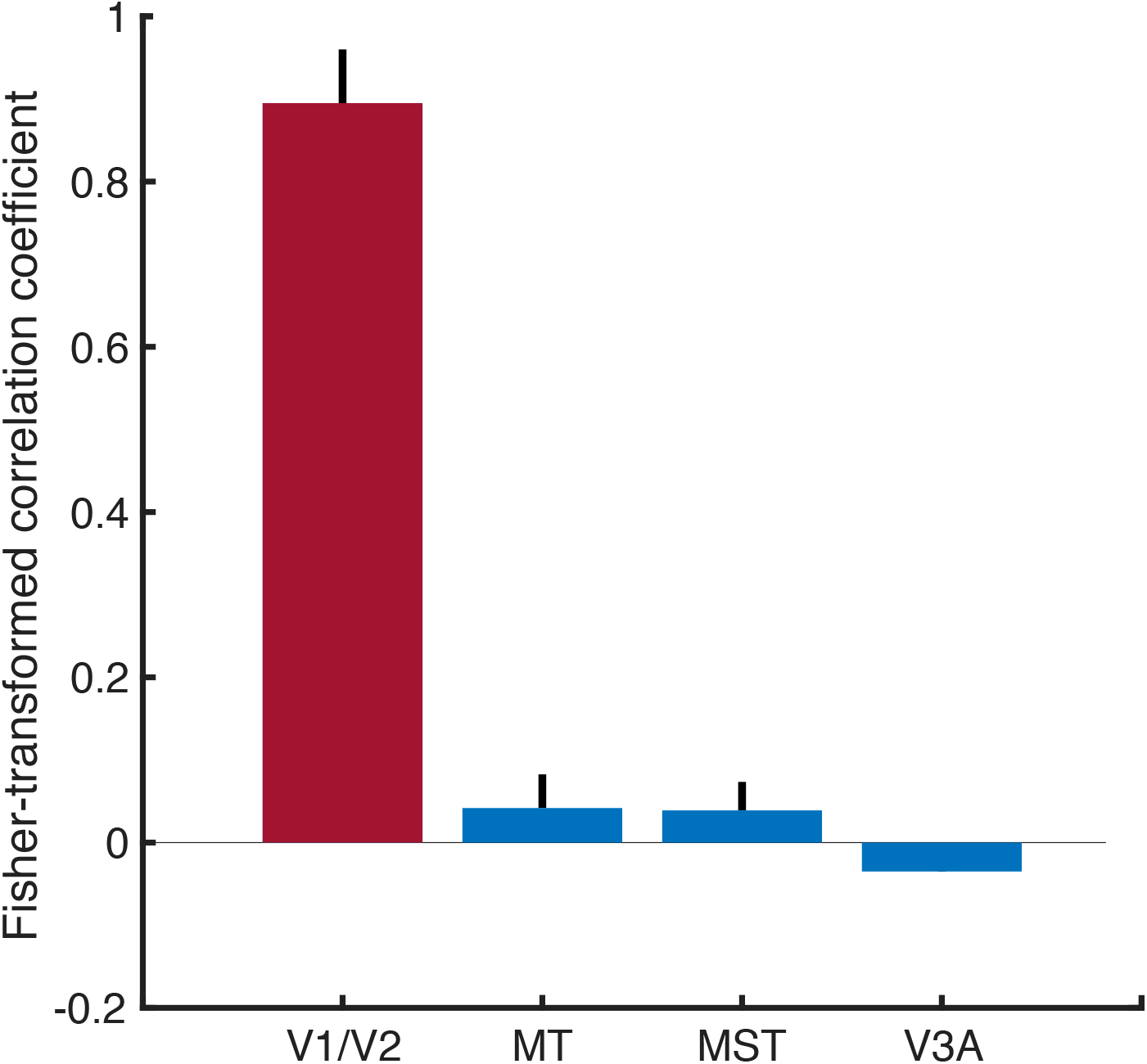
Mean Fisher-transformed correlation coefficient between the estimated scores of the V1/V2 activation patterns during the neurofeedback stage and the reconstructed scores of the activation patterns in V1/V2 (red) and in other control ROIs (blue). Only the activation pattern in V1/V2 (red) can significantly reconstruct the estimated scores of V1/V2 activation pattern during neurofeedback stage.

### Control Experiment

To rule out that the improvement in the local motion ranges were induced by performing motion discrimination tasks only during pre- and posttest stages or by exposure during the decoder construction stage, we conducted a control experiment (see **Materials and Methods** for detail). Subjects performed the same task in the testing stages as in the neurofeedback experiment. During the decoder exposure stage, the same stimulus used during the decoder construction stage was presented to the subjects outside the scanner. Since there was no trained versus untrained local motion ranges, we specified the two types of global motion display as clockwise or counterclockwise of vertical. We did not find any significant performance improvement in either clockwise [t(5) = 0.03, p = 0.9771] or counterclockwise local motion ranges [t(5) = −0.487, p = 0.647] (shown in Figure 7), unlike in the neurofeedback experiments.

**Figure 7.**
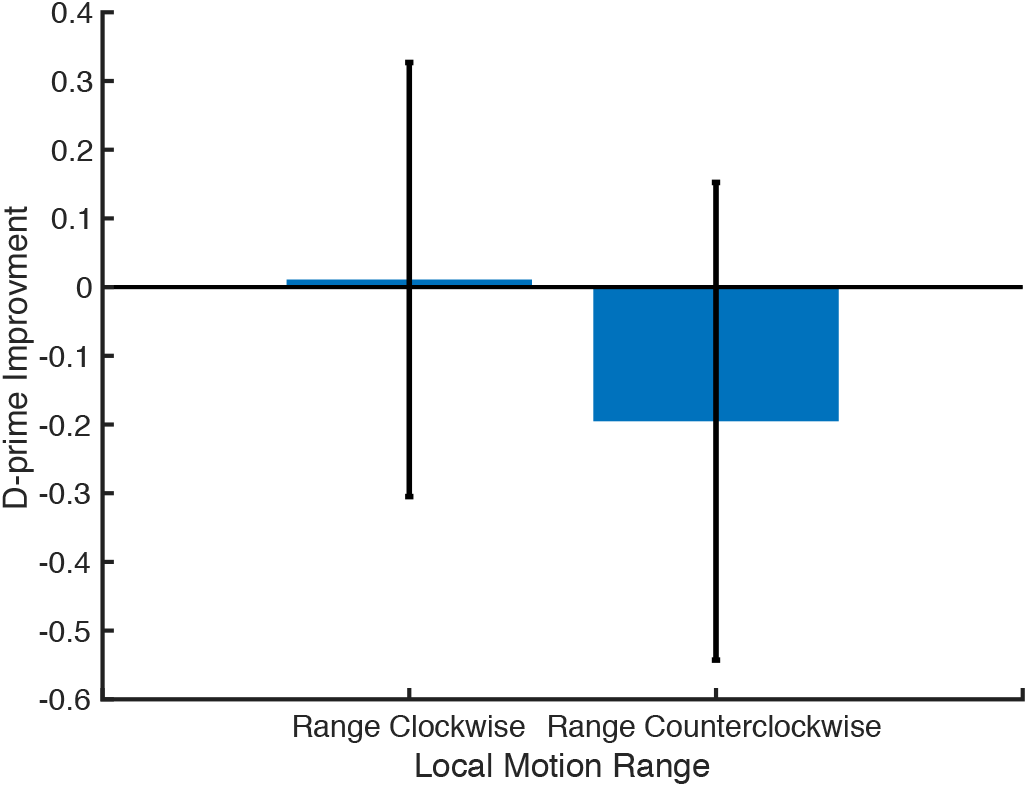
Local motion range improvement in the control experiment. No significant change was observed for the two local motion ranges.

## Discussion

A psychophysical study (Watanabe et al., 2002) found that exposure to a global motion display improved the sensitivity to the local motion direction range but not specifically the global motion direction. The early phases of training on global motion discrimination in the global motion display led to performance improvement in the local motion direction ranges, whereas in later phases of training, performance improvement occurred in the global motion direction (Watanabe et al., 2002). These results suggest that exposure to a local feature leads to improvement in the sensitivity to the feature, irrespective of whether the feature is task relevant. However, these psychophysical-only results left us with at least two questions. First, what cortical area or areas are involved in this exposure-based plasticity? Second, does exposure-based plasticity truly occur passively? The fact that local motion directions were task-irrelevant does not unequivocally indicate that attention was not directed to those directions.

In the current study, we trained subjects with online decoded neurofeedback in which a decoded global motion pattern was repetitively induced in V1/V2 while the subjects were not presented with a global motion display and were not aware of the purpose of the experiment. As a result, performance improved only in the local motion ranges and not specifically in the global motion direction. Offline leak analyses confirmed that the activation pattern related to feedback during neurofeedback training was largely confined to V1/V2. These results are in accordance with the hypothesis that V1/V2 is involved in exposure-based plasticity in VPL, which occurs passively and in an unconscious manner without involving attention from higher areas.

Several of our results consistently suggest that the behavioral improvement developed with DecNef is restricted to neural activity and, in particular, neural representation in the target region. First, DecNef training reduces or eliminates the possibility that modified functions after neurofeedback training are ascribed to neural pattern changes involved in subjects’ intentions to improve the functions (Shibata et al., 2019; Watanabe et al., 2017). While the subjects were aware of the feedback signals themselves, they were not aware of either what was trained or even the fact that they were being trained. Verbal reports after the experiment (see Table 1) indicate that the subjects’ intended strategies (see **Materials and Methods** for details) during the neurofeedback stage did not include anything to do with motion. Second, the improvement was confined to the local motion range. As discussed above, V1 is the earliest human cortical area that responds to local motions, while V3A is the earliest area that responds to global motion (Koyama et al., 2005). The improvement in the local motion directions and not specifically in the global motion direction suggests that DecNef training is effective on areas lower than V3A. Third, the results of the leak analysis indicate that the activation pattern related to neurofeedback during neurofeedback training was confined to V1/V2. The leak analysis of the activation patterns during the neurofeedback training stage throughout the brain showed that the activity patterns representing motion information similar to that in V1/V2 were confined to V1/V2 instead of other motion-related brain regions.

Is it plausible to conclude that exposure-based VPL can be causally driven by changes in V1/V2? In order to gain an understanding of the causal relationship between an activity pattern in a certain brain area and a certain behavioral function, it is necessary for a real-time neurofeedback technique to reliably activate the targeted pattern in the targeted area and to specifically change the targeted behavioral function. The results of the current and previous studies suggest that it is reasonable to suggest a causal relationship between a brain activation pattern and a corresponding behavioral function. First, these DecNef studies show that the targeted activity patterns in a targeted area are unlikely to spread out to other areas by means of leak analysis (Shibata et al, 2010; Amano et al, 2016; Shibata et al, 2016). Second, the current study demonstrated that a specific behavioral function is changed in an aimed fashion. As discussed above, the global motion display used for the current DecNef training, if physically presented, induces the perception of both local motions and the global motion in the direction of the spatiotemporal average of the local motion vectors. However, the improvement in the sensitivity only to the local motion directions due to the current DecNef training indicates that DecNef specifically changes a targeted behavior function. Thus, our study suggests a causal relationship between V1/V2 and exposure-based VPL.

In a two-stage model of VPL (Shibata, Sagi, & Watanabe, 2014; Watanabe et al., 2002; Watanabe & Sasaki, 2015), VPL of a feature consists of feature-based plasticity, which refers to the changes in a feature representation as a result of exposure, and of task-based plasticity, which refers to the changes in high-level stages involving task processing. The results of the current DecNef experiment support the feature-based plasticity aspect of the model in at least the following ways. First, the results indicate that VPL of local motion can occur without the involvement of high-level processing during training that was driven by the subjects’ clear intention of improving sensitivity to the local motions. Second, the results also show that passive training led to changes only in low-level stages such as V1/V2. Early visual areas, including V1/V2, have been thought to be heavily involved in processing related to a feature representation (Hubel & Wiesel, 1963).

In summary, activity patterns in V1/V2 similar to those evoked by a real presentation of a global motion display were induced by DecNef training without the actual presentation of a trained stimulus, leading to sensitivity increases to local motion directions but not to the global motion direction, even though the actual presentation of the global motion display would induce the perception of both types of motions. This result is in accordance with the hypothesis that VPL of a low-level visual feature can occur in V1/V2 in a passive and unconscious manner. (981/1500 words)

## Author contributions

ZW, YS and TW designed the study. ZW and MT performed the experiments. KS, MW and TY provided technical support for real-time neurofeedback. ZW analyzed the data. ZW, KS, TY, MK, YS and TW wrote the manuscript.

## Acknowledgements

Z.W., T.Y., Y.S. and T.W. and the study was supported by NIH R21EY028329, R01EY027841, R01EY019466, and United States - Israel Binational Science Foundation BSF2016058. Z.W. was also supported by T32MH115895. M.K. was partially supported by AMED (Japan, JP18dm0307008). K.S. was partially supported by JSPS Kakenhi (19H01041). M.S.W was supported by NIGMS-NIH P20GM103645.

